# Identification of 67 pleiotropic genes for seven autoimmune diseases using multivariate statistical analysis

**DOI:** 10.1101/563973

**Authors:** Xiaocan Jia, Nian Shi, Zhenhua Xia, Yu Feng, Yifan Li, Jiebing Tan, Fei Xu, Wei Wang, Changqing Sun, Hongwen Deng, Yongli Yang, Xuezhong Shi

## Abstract

Although genome-wide association studies (GWAS) have a dramatic impact on susceptibility locus discovery, this univariate approach has limitation in detecting complex genotype-phenotype correlations. It is essential to identify shared genetic risk factors acting through common biological mechanisms of autoimmune diseases with a multivariate analysis. In this study, the GWAS summary statistics including 41,274 single nucleotide polymorphisms (SNPs) located in 11,516 gene regions was analyzed to identify shared variants of seven autoimmune diseases using metaCCA method. Gene-based association analysis was used to refine the pleiotropic genes. In addition, GO term enrichment analysis and protein-protein interaction network analysis were applied to explore the potential biological function of the identified genes. After metaCCA analysis, 4,962 SNPs (*P*<1.21×10^−6^) and 1,044 pleotropic genes (*P*<4.34×10^−6^) were identified. By screening the results of gene-based p-values, we identified the existence of 27 confirmed pleiotropic genes and highlighted 40 novel pleiotropic genes which achieved significance threshold in metaCCA analysis and were also associated with at least one autoimmune disease in the VEGAS2 analysis. The metaCCA method could identify novel variants associated with complex diseases incorporating different GWAS datasets. Our analysis may provide insights for some common therapeutic approaches of autoimmune diseases based on the pleiotropic genes and common mechanisms identified.

**Author summary:** Although previous researches have clearly indicated varying degrees of overlapping genetic sensitivities in autoimmune diseases, it has proven GWAS only explain small percent of heritability. Here, we take advantage of recent technical and methodological advances to identify pleiotropic genes that act on common biological mechanisms and the overlapping pathophysiological pathways of autoimmune diseases. After selection using multivariate analysis and verification using gene-based analyses, we successfully identified a total of 67 pleiotropic genes and performed the functional term enrichment analysis. In particularly, 27 genes were identified to be pleiotropic in previous different types of studies, which were validated by our present study. Forty significant genes (16 genes were associated with one disease earlier, and 24 were novel) might be the novel pleiotropic candidate genes for seven autoimmune diseases. The improved detection not only yielded the shared genetic components but also provided better understanding for exploring the potential common biological pathogenesis of these major autoimmune diseases.

## Introduction

Autoimmune diseases are chronic conditions initiated by loss of immunological tolerance to self-antigens[1]. An estimated incidence of autoimmune diseases is about 90 cases per 100,000 person-year and the prevalence is about 7.6–9.4% in Europe and North America[2]. The chronic nature of such diseases has a significant impact in terms of the utilization of medical care, direct and indirect economic costs and quality of life. In addition, extensive clinical and epidemiologic observations have shown that autoimmune diseases are characterized by familial clustering of multiple diseases, epidemiological co-occurrence, and the efficacy of therapies across diseases, which means different autoimmune diseases share a substantial portion of their pathobiology and genetic predisposition underlies disease susceptibility[3, 4].

Pleiotropy describes the genetic effect of a single nucleotide polymorphism (SNP) or gene on two or more phenotypic traits and its outcome is genetic correlation. Largely, this concept concerns across-trait architecture[5]. Recent genome wide association studies (GWAS) in autoimmune diseases and subsequent replication studies have identified 186 susceptibility loci with statistically significant and more than half of them are shared by at least two distinct autoimmune diseases[6–9]. For example, *PTPN22* c.1858C>T (rs2476601), is evident in independent GWAS across multiple autoimmune diseases[10]. In addition, there is evidence that loci predisposing to one disease can have effects on risk of a second disease, although the risk allele for one disease may not be the same as for the second[6]. However, evidence for specific shared risk variants is modest, and consequently the genetic mechanisms that may explain the patterns of disease aggregation remain unclear. It is therefore important to identify more shared genetic risk factors acting through common biological mechanisms and assess overlapping pathophysiological relationships of autoimmune diseases using effective analytical approaches.

GWAS is a standard univariate approach to investigate and identify potentially causal or risk-conferring genetic variants for common human diseases in the individual level measurement[11, 12]. GWAS, especially those with large sample size and meta-analysis of multiple studies, have a dramatic impact on susceptibility locus discovery and in addition, highlight and extend the previously observed commonality of loci between autoimmune diseases. However, with millions of SNPs and a growing number of phenotypes, this univariate approach has had limited success in detecting complex genotype-phenotype correlations. Furthermore, the researches of statistical methods have proved multivariate analysis had higher statistical power to detect the unexplained heritability due to considering correlations not only among multiple SNPs but also among different traits or diseases[13]. Existing studies of genetic risk factors for complex traits have used bivariate analysis, but multivariate analysis based on complex diseases is rare[14, 15]. Therefore, a multivariate analysis to identify pleiotropic genes, especially using the publicly available summary statistics of GWAS, is very essential and relevant.

Cichonska et al[16] recently performed a canonical correlation analysis method allowing multivariate representation of both genotypic and phenotypic variables based on the published univariate summary statistics from GWAS by meta-analysis(metaCCA). This new approach has been applied to identify potential pleiotropic genes in lipid related measures, psychiatric disorders, and chronic diseases[16–19]. In this report, the genetic pleiotropy-informed metaCCA method was used to identify shared variants and pleiotropic effect in seven autoimmune diseases: celiac disease (CEL), inflammatory bowel disease (IBD, which includes Crohn’s disease (CRO) and ulcerative colitis (UC)), multiple sclerosis (MS), primary biliary cirrhosis (PBC), rheumatoid arthritis (RA), systemic lupus erythematosus (SLE) and type 1 diabetes (T1D). In addition, gene-based association analysis was used to refine pleiotropic genes. GO term enrichment analysis and protein-protein interaction network analysis were applied to explore the potential biological function of the identified genes.

## Results

### Potential pleiotropic SNPs and genes by metaCCA analysis

After gene annotation and SNP pruning, there were 41,274 SNPs located in 11,516 gene regions available for the metaCCA analysis. The size of SNP representation of the genes ranged from 1 to 213 SNPs, and the median number of SNPs in each gene was 3.72. For the univariate SNP-multivariate phenotype analysis, 4,962 SNPs reached the Bonferroni corrected threshold (p<1.21×10^−6^), and the canonical correlation r between each SNP and phenotype ranged from 0.0372 to 0.6586. The results are presented by the Manhattan plot in **Fig. 1**. For the multivariate SNP-multivariate phenotype analysis, 1,044 genes with a significance threshold of p<4.34×10^−6^ were identified as the potential pleiotropic genes. The canonical correlation r between genotype and phenotype ranged from 0.0322 to 0.5899.

**Figure 1.**
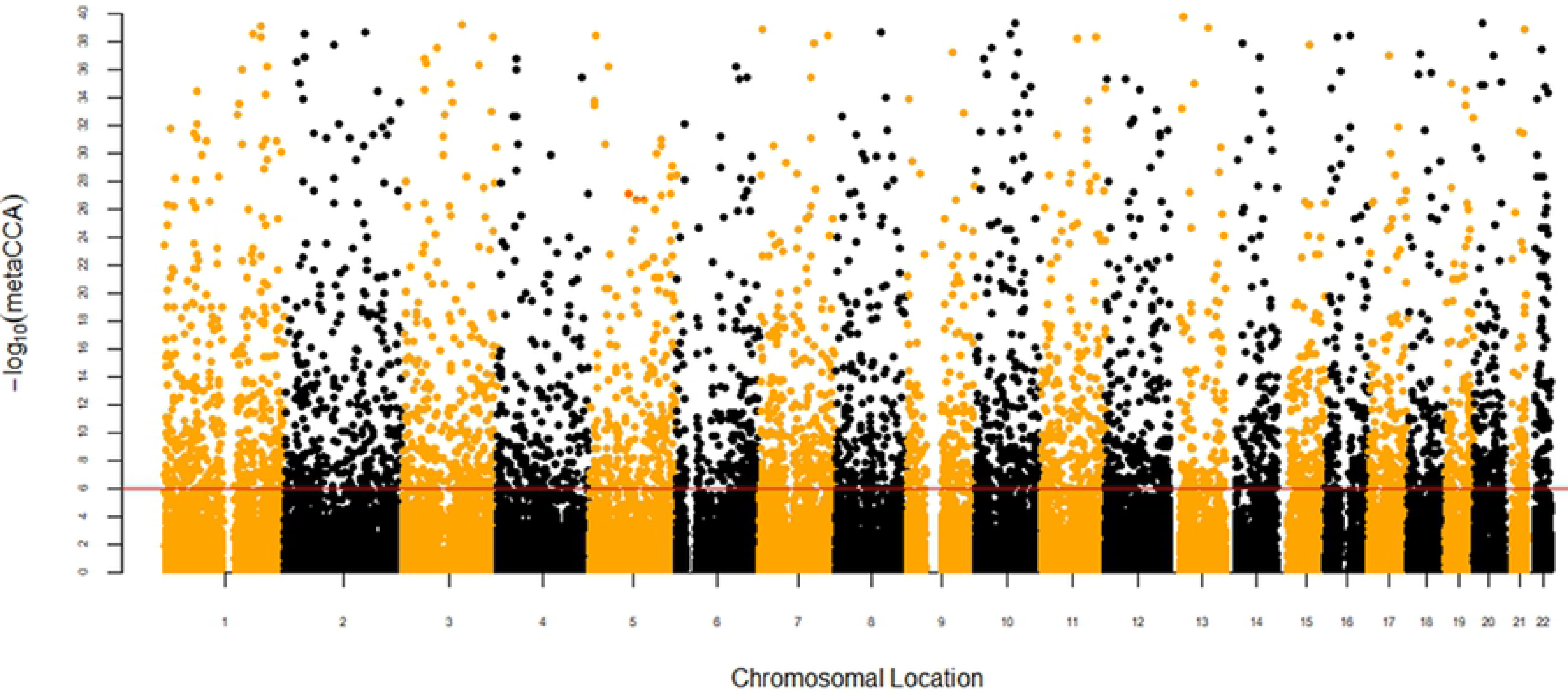
Manhattan plot of −log_10_(metaCCA) values for univariate SNP-seven autoimmune diseases analysis. The red line marks the −log_10_(metaCCA) value of 5.92 corresponding to p<1.21×10^−6^. If the −log_10_(metaCCA) value of a certain SNP was greater than 5.92, this SNP was identified as a pleiotropic SNP for seven autoimmune diseases.

### Refining the pleiotropic genes by gene-based association analysis

VEGAS2 algorithm based on original GWAS summary statistics was used to identify association of one gene with specific disease. After the gene-based association analysis, 19 genes were identified for CEL, 111 genes were identified for IBD, 16 genes were identified for MS, 20 genes were identified for PBC, 19 genes were identified for RA, 20 genes were identified for SLE, and 33 significant genes were identified for T1D with the p-value equal to 1.00E-06.

By screening the results of gene-based analysis p-values, we identified 67 putative pleiotropic yielding significance in the metaCCA analysis and were associated with at least one disease in the VEGAS2 analysis. In particular, 17 genes were associated with more than one disease in the original GWAS. The findings of the metaCCA and VEGAS2 analysis are summarized in Specifically, 27 of these 67 putative pleiotropic genes had been previously reported to be associated with more than one of these seven diseases. Of these 27 confirmed pleiotropic genes, 6 genes (*ADAD1*, *CIITA*, *CLEC16A*, *IL23R*, *MAGI3*, *PTPN2*) were associated with more than one disease in the VEGAS2 analysis using original GWAS summary statistics. Of the 40 detected novel putative pleiotropic genes, 16 genes were previously reported to be associated with only one autoimmune disease. *EFR3B* and *RBM17* were reported to be only associated with T1D in published studies, but were associated with multiple diseases in the VEGAS2 analysis. Other 24 remaining significant genes might represent candidate novel pleiotropic genes for these autoimmune diseases. More significantly, 9 genes (*C1orf141*, *CALU*, *CCDC136*, *FGF2*, *LOC101927051*, *MAP4K4*, *MPZL3*, *PAPOLG*, *SH3BP1*) were associated with more than one disease in the VEGAS2 analysis although they had never been reported to be associated with any autoimmune disease. The detailed features of 67 significant pleiotropic genes are shown in **Table 2.**

**Table 1.**
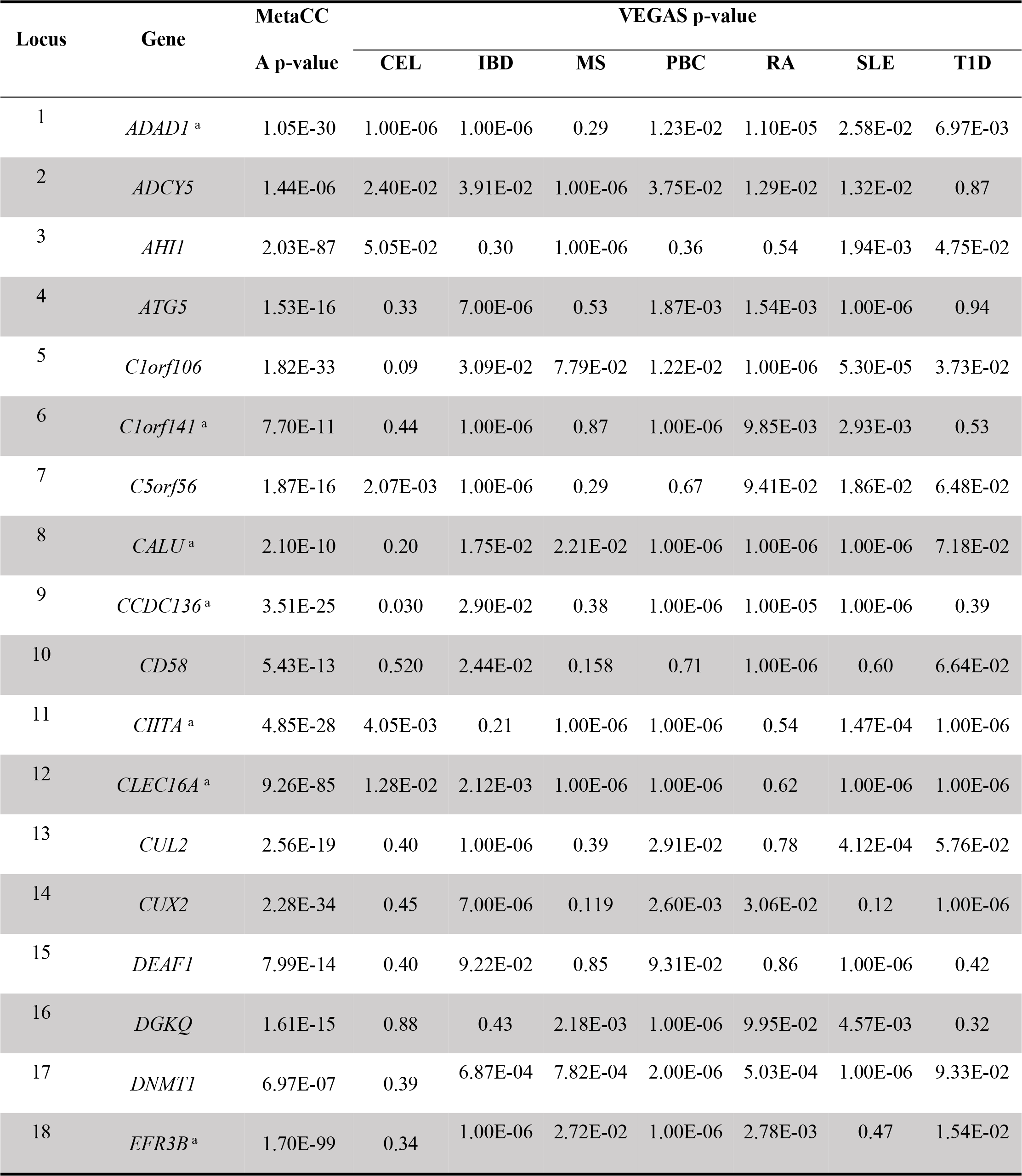
The 67 pleiotropic genes identified by the metaCCA and VEGAS2 analysis

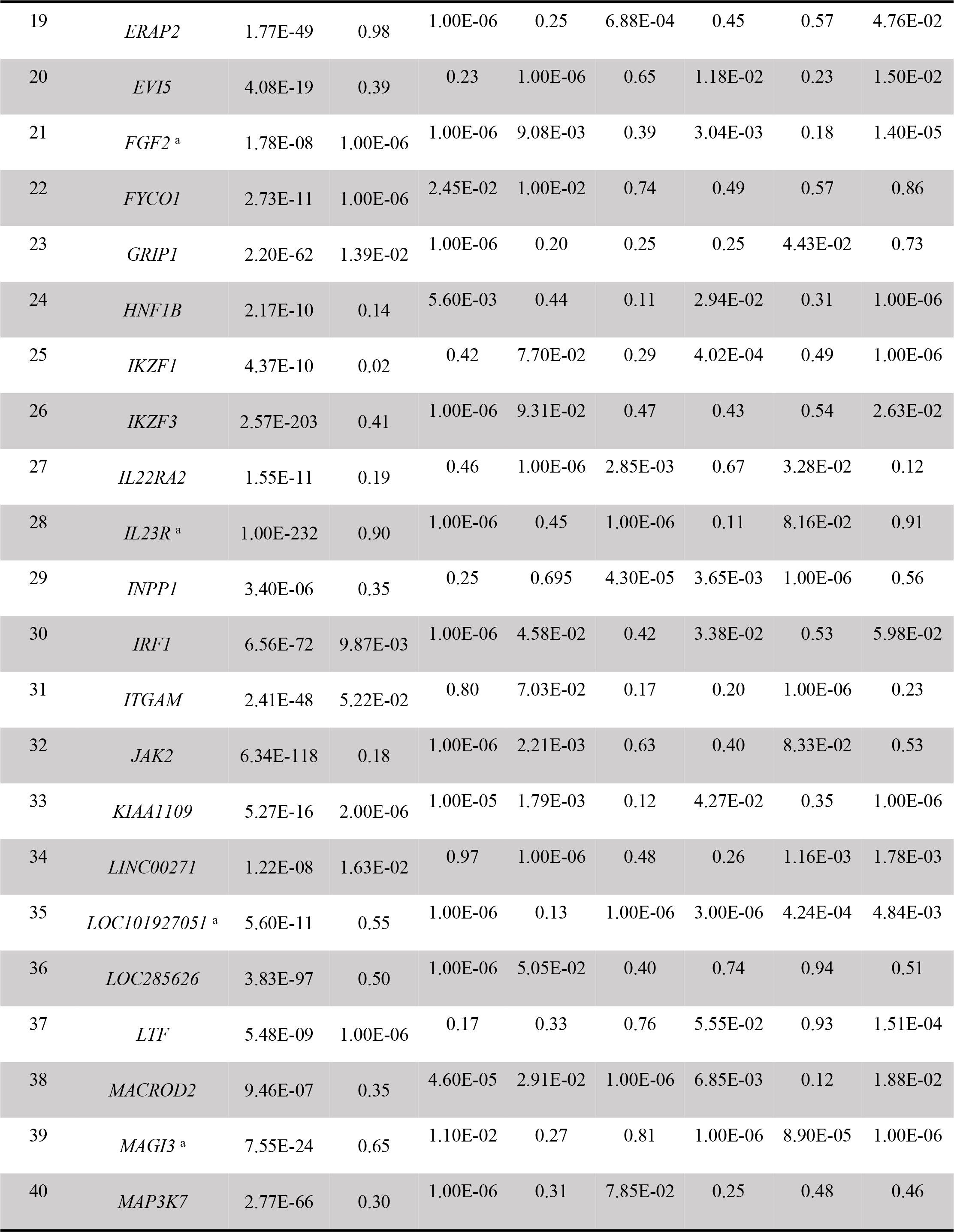

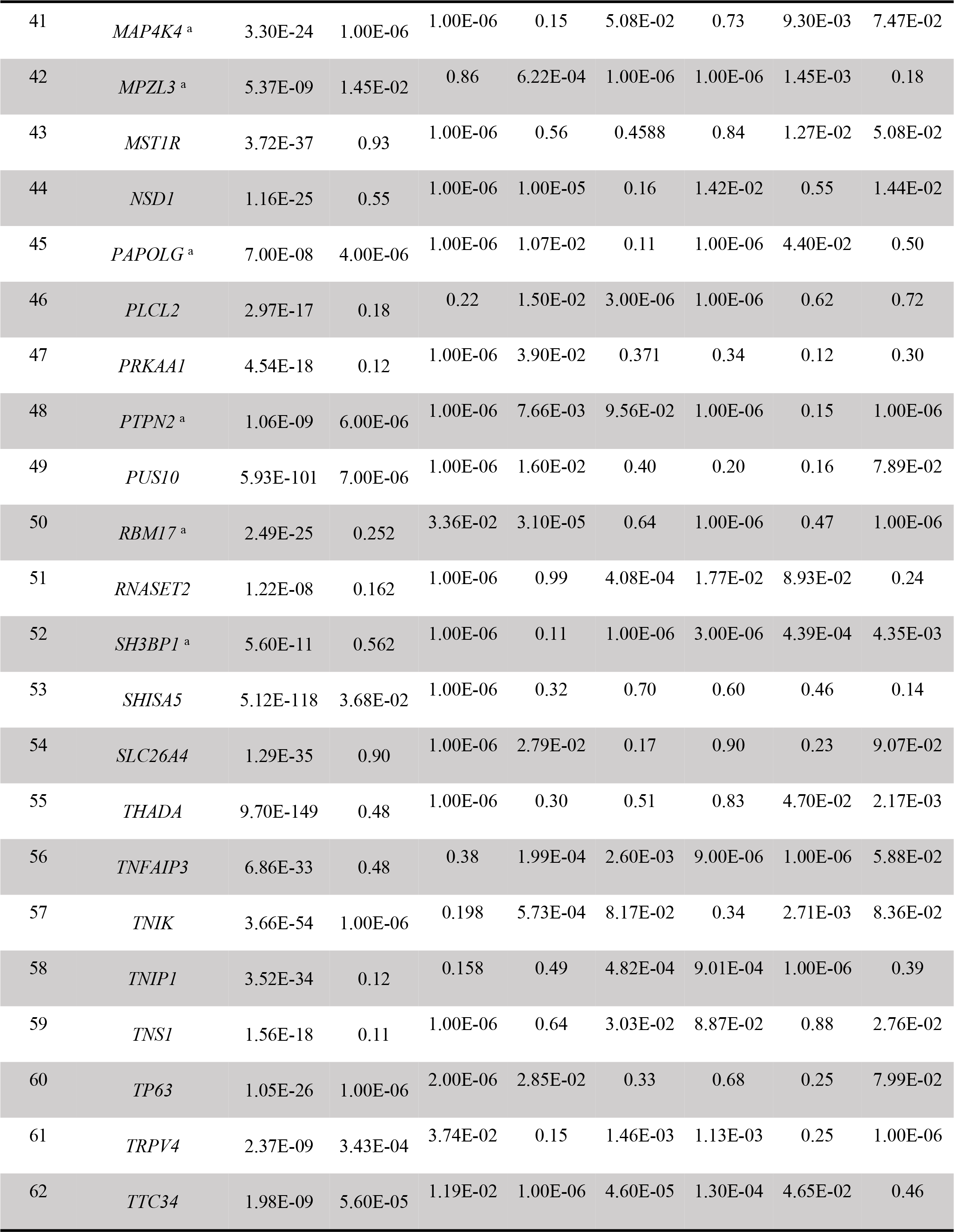

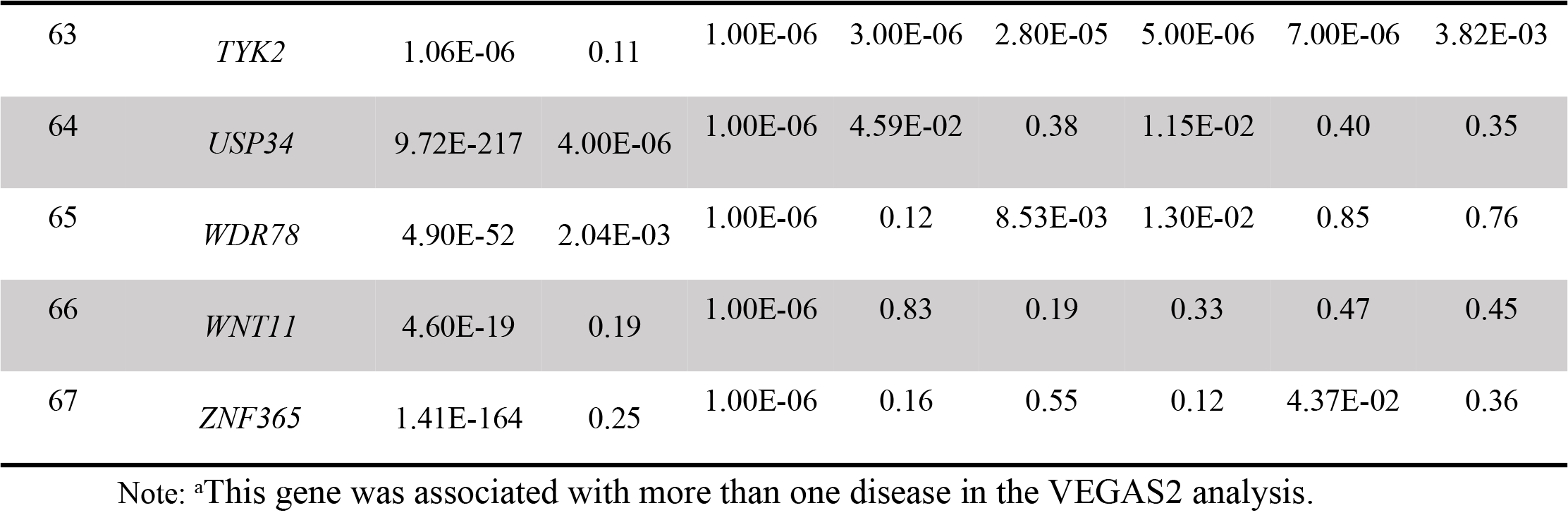

**Table 2.**
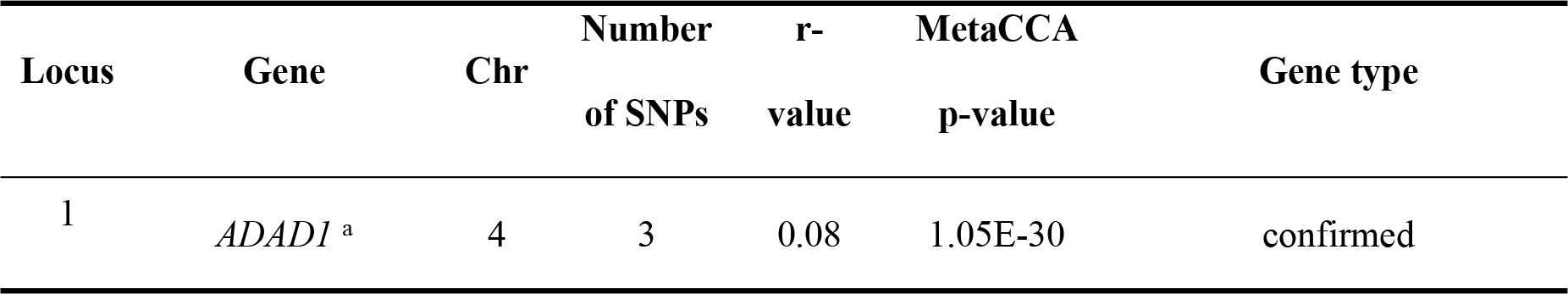
The features of 67 significant pleiotropic genes

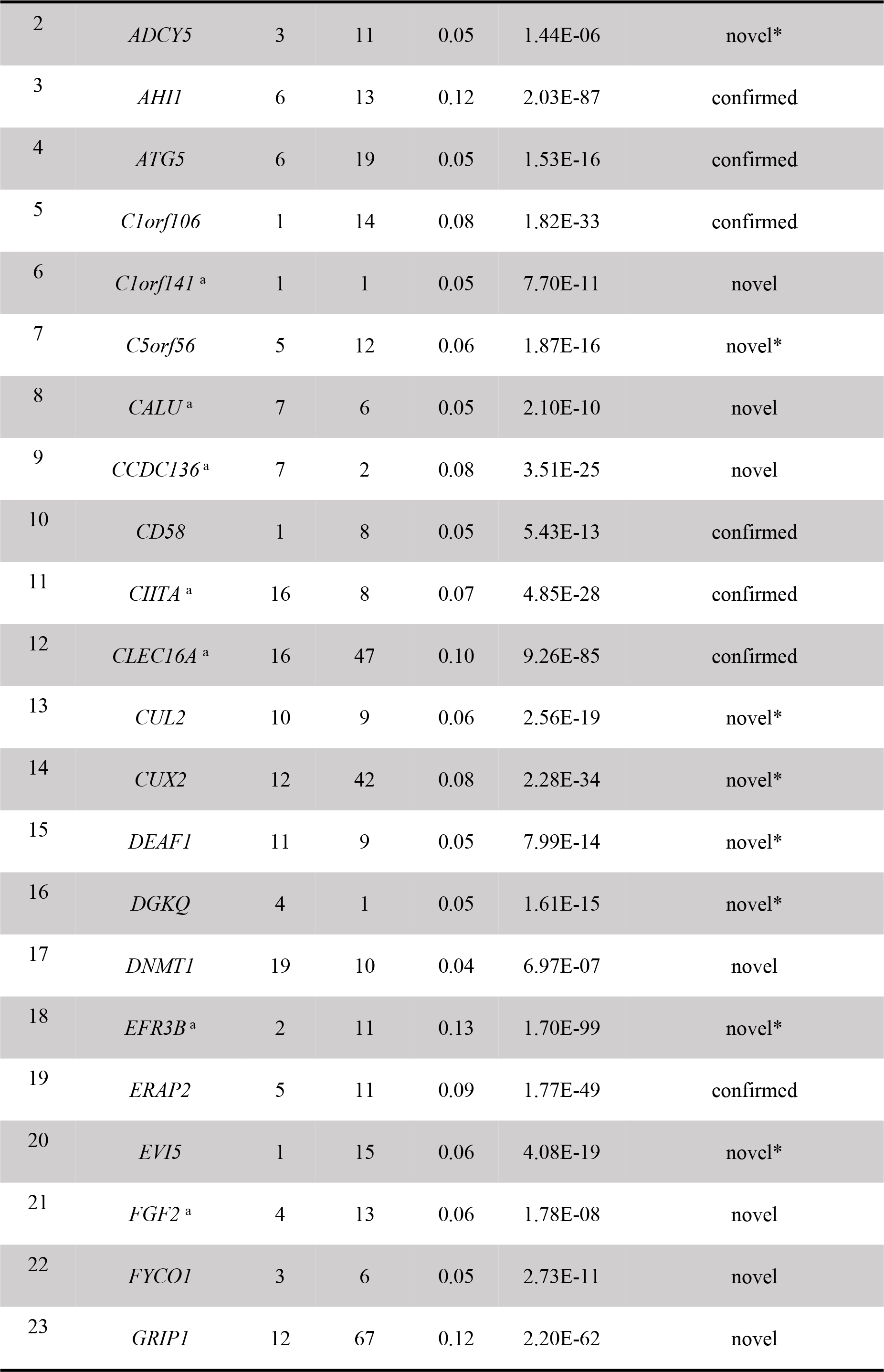

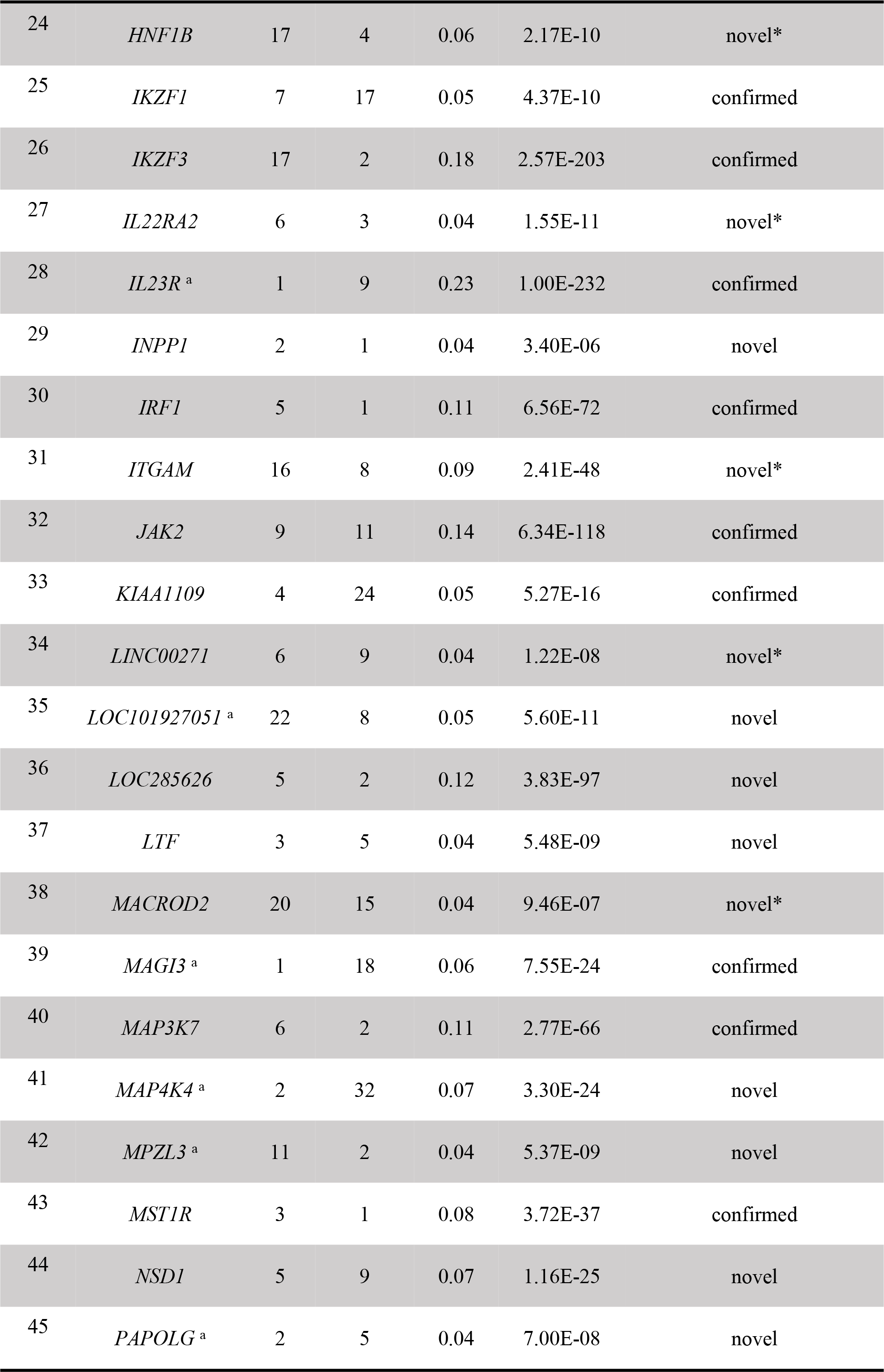

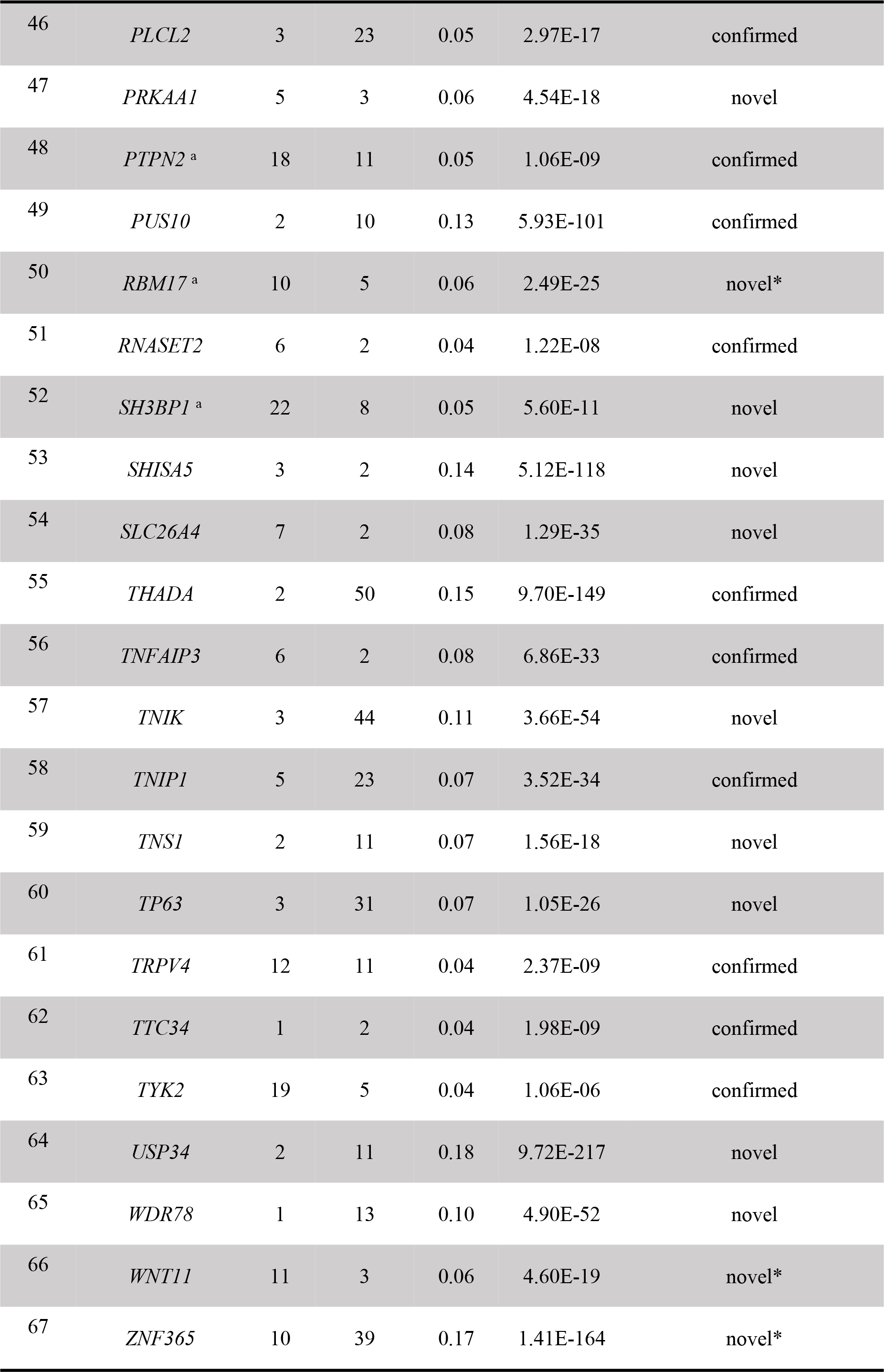

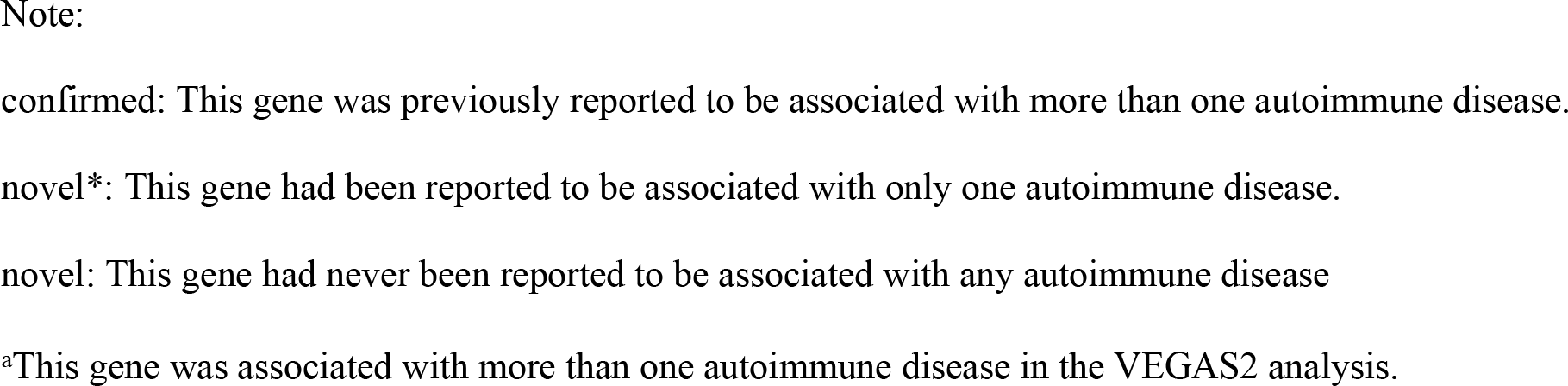

### GO term enrichment analysis

GO enrichment analysis revealed that the biological functions of these pleiotropic genes were mainly involved in the metabolism of lipids. When 67 pleiotropic genes associated with autoimmune diseases were used as the gene sets for the GO term enrichment analysis, several functional terms were identified as being enriched. For the GO biological process and molecular function, there were 63 and 5 significant terms were identified to be enriched in the development of autoimmune diseases, respectively. The detailed information of top five significant GO terms is shown in **Table 3**.

**Table 3.**
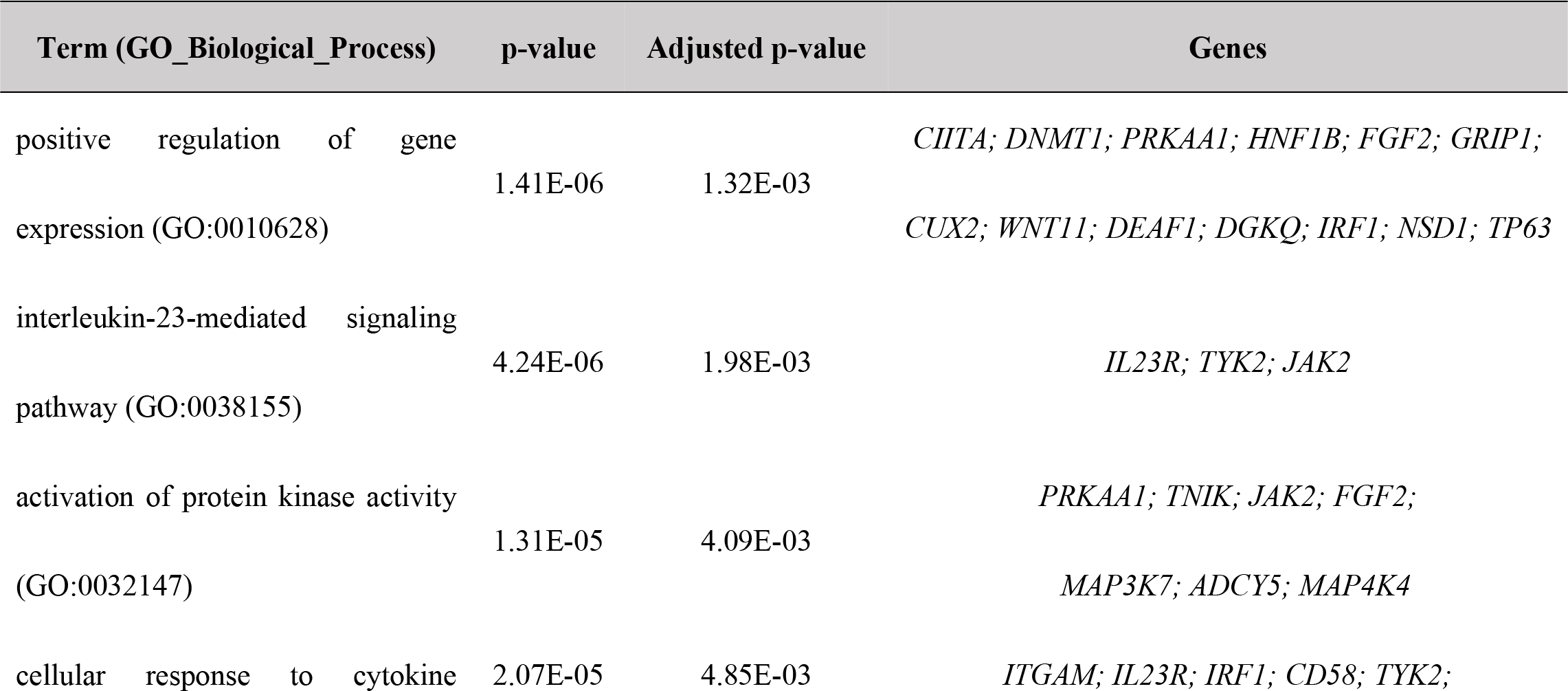
Top five significant GO Term enrichment of the 67 pleiotropic genes

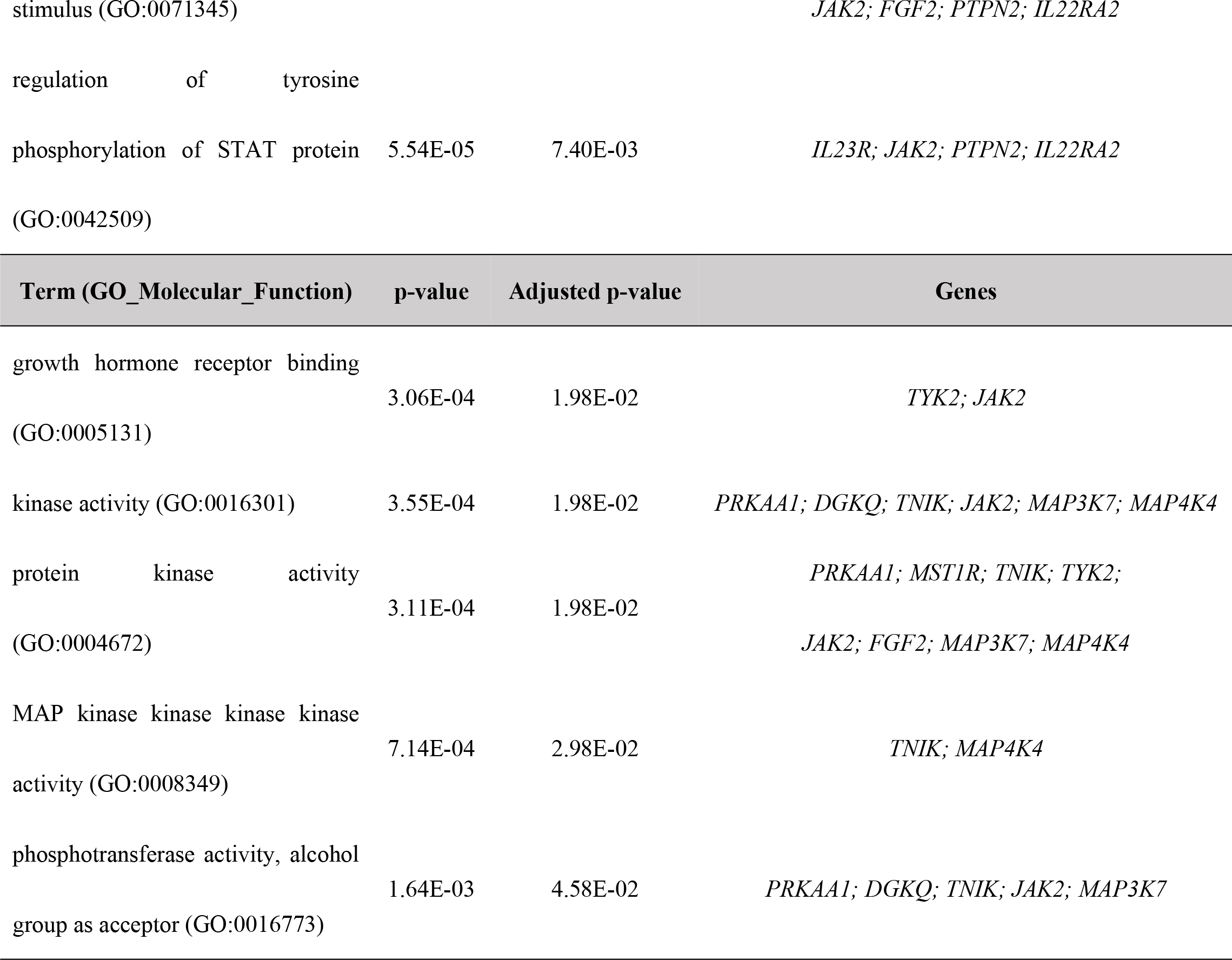

### Protein-protein interaction network analysis in STRING

The 67 identified putative pleiotropic genes were retrieved from the STRING database. The 63 genes were clearly enriched in two confirmed clusters: *JAK2* and *MAP3K7* clusters (**Fig. 2**). Three detected novel putative pleiotropic genes, *FGF2*, *IL22RA2*, and *ITGAM* were involved in the *JAK2* cluster, which participates in various processes such as cell growth, development, differentiation or histone modifications and mediates essential signaling events in both innate and adaptive immunity. Three other novel genes, *MAP4K4*, *PRKAA1* and *TNIK* were involved in the *MAP3K7* cluster, which acts as an essential component of the MAP kinase signal transduction pathway and plays a role in the response to environmental stress and cytokines such as TNF-alpha.

**Figure 2.**
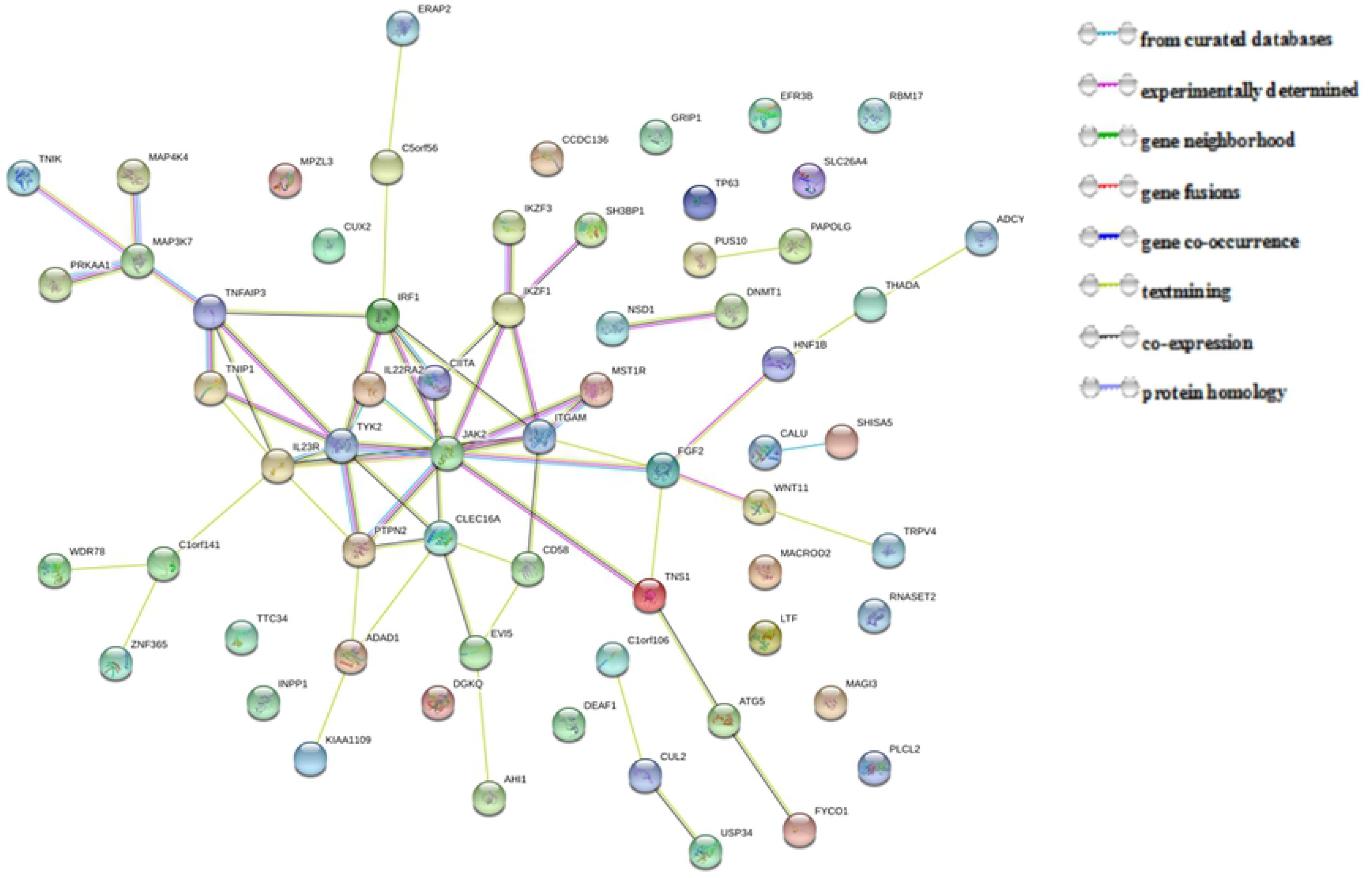
Protein-protein interactions between 67 pleiotropic genes associated with autoimmune diseases.

## Discussion

In the present study, a novel analytical approach – metaCCA was used to explore the common genetic variants for autoimmune diseases by combining seven available independent GWAS or meta-analysis summary statistics. A total of 67 putative pleiotropic genes were successfully identified after verification using gene-based analysis. In particularly, 27 confirmed genes were identified as pleiotropic in previous different types of studies and were validated in the present study, 16 novel pleiotropic genes were previously reported to be associated with one autoimmune disease, and 24 candidate novel pleiotropic genes had never been reported to be associated with any autoimmune disease. The improved detection and biological pathways may provide with novel insights into the shared genetic factors in development of autoimmune diseases.

Among the 27 confirmed pleiotropic genes, 6 genes (*ADAD1*, *CIITA*, *CLEC16A*, *IL2*3R, *MAGI3*, *PTPN2*), which play an important role on the pathomechanism of autoimmune diseases, were proven to be associated with more than one autoimmune disease not only in literature review but also in the VEGAS2 analysis using original GWAS summary statistics. For example, common genetic variants in *CLEC16A*, also called the C-type lectin-like domain family 16A, had been reported to be associated with CEL, IBD, MS, PBC and T1D[7]. As the non-HLA genome-wide significant risk gene, *CLEC16A* is essential for autophagosomal growth and autophagy processes, which are of major importance for proper immune regulation, including regulation of inflammasome activation[20, 21]. Most of all, recent murine data and protein-protein interaction network analysis in our study indicated that *CLEC16A* plays a key role in beta cells functions by regulating mitophagy/autophagy and mitochondrial health[22]. *PTPN2* is another important and confirmed pleiotropic gene associated with several autoimmune diseases we studied[7]. The GO term enrichment analysis results suggest *PTPN2* encodes the T-cell protein tyrosine phosphatase, acting as a negative regulator of the JAK/STAT signaling pathways downstream of cytokines and playing a prominent role in T-cell activation, signaling and/or effector function, which may be the potential targets for the pharmacotherapy of autoimmune diseases. In addition, Mei QB et al[23] also shown that *PTPN2* genetic polymorphisms are associated with psoriasis in the Northeastern Chinese population, and psoriasis is another chronic immune-mediated disease with a complex etiology.

Sixteen detected novel putative pleiotropic genes had been early validated associated with one kind of autoimmune disease. Interestingly, *EFR3B* and *RBM17* had been reported to be only associated with T1D in published studies, but were validated associated with other diseases in the VEGAS2 analysis[10, 24, 25]. *EFR3B* is an associated gene located in 2p23, which probably acts as the membrane-anchoring component and involved in responsiveness to G-protein-coupled receptors. Although Bradfield et al[10] have validated *EFR3B* is an associated loci and protein-protein interaction network analysis also provide some protein information, it is still unclear whether this role is direct or indirect. *RBM17*, involved in the regulation of alternative splicing and the utilization of cryptic splice sites, is essential for survival and cell maintenance. Fortunately, genetic and serologic data suggest that the inherited altered genetic constitution located between *IL2RA* and *RBM17* may predispose to a less destructive course of RA[26]. Although the 14 remaining genes were identified associated with one kind of autoimmune disease in literature review and the VEGAS2 analysis, different experimental study may provide sufficient evidence to support them as pleiotropic genes for autoimmune diseases. *CUL2* is an associated gene with response to the hypoxic environment and activation of tumor immune system, which has been identified associated with CRO in nine independent case-control series[27]. Zhang, WY et al[28] suggest that human immunodeficiency virus type 1 and simian immunodeficiency virus viral infectivity factor form a CRL5 E3 ubiquitin ligase complex to suppress virus restriction by host APOBEC3 proteins, and eventually *CUL2* may suppress this pathway and increase the risk of autoimmune diseases[29].

Significantly, 9 genes (*C1orf141*, *CALU*, *CCDC136*, *FGF2*, *LOC101927051*, *MAP4K4*, *MPZL3*, *PAPOLG*, *SH3BP1*) were associated with more than one disease in the VEGAS2 analysis although they had never been reported to be associated with any autoimmune disease in previous GWAS. *MAP4K4* has been enriched several GO terms including MAP kinase kinase kinase kinase activity (GO:0008349), and this term act as an important contributor to the risk toward the development of Type 2 Diabetes Mellitus in a Chinese Han Population[30]. In addition, Aouadi, M et al[31] demonstrated that Orally delivered siRNA targeting macrophage *MAP4K4* suppresses systemic inflammation. This technology defines a new strategy for oral delivery of siRNA to attenuate inflammatory responses in human disease. *C1orf141* is another significant candidate novel pleiotropic gene testified associated with IBD and PBC in our study. It has been recently shown that *C1orf141* is a susceptibility variant of psoriasis, a chronic inflammatory hyperproliferative cutaneous disease[32]. Here, we don’t have a detailed description of each candidate novel gene because pathomechanisms are unclear apparently, and further experimental studies will need to be conducted to confirm our novel findings.

Systematically and comprehensively searching for the pleiotropic genes and their effects is essential and necessary for understanding development mechanisms of autoimmune diseases[3]. Compared to the univariate GWAS analysis based on cross-sectional population, it was a cost-effective and reliable study, which not only increased sample size by integrating seven large GWAS summary statistics, but also increased statistical power and provided richer findings by jointly analyzing multiple related autoimmune diseases. However, this study could not relate to the information about the direction of effects of pleiotropic genes on risk to these diseases because of a lack of detailed original individual measures. Alternative approaches and experimental studies may be applied to check whether novel genes could still be identified/substantiated with these methods in order to confirm novel findings in the further study.

In summary, we provided convincing evidence for the existence of 27 confirmed pleiotropic genes and highlighted 40 novel pleiotropic genes for autoimmune diseases by performing a systematic multivariate analysis of the open GWAS data using metaCCA. Furthermore, we illustrated potential biological functions of this pleiotropic genes and our results can contribute to a better understanding of common genetic mechanisms, and eventually the development of improved diagnosis, prognosis and targeted therapies.

## Materials and Methods

### GWAS Datasets

All the GWAS summary statistics of seven autoimmune diseases in this present study were downloaded from ImmunoBase (website: https://www.immunobase.org/), which is a web based resource focused on the genetics and genomics of immunologically related human diseases. The data of CEL was a second-generation GWAS of 4,533 cases and 10,750 control subjects including 523,402 SNPs[33]. The association summary statistics of IBD including 9,735,446 imputed SNPs was obtained from a meta-analysis with a total sample size of 59,957 subjects[34]. The dataset for MS was 464,357 genotyped or imputed SNPs from a collaborative GWAS involving 9772 cases and 17,376 controls of European descent collected by 23 research teams from 15 different countries[35]. The dataset for PBC including 1,134,141 SNPs was from a meta-analysis (2,764 cases and 10,475controls) and an independent cohort (3,716 cases and 4,261 controls)[36]. RA was also a GWAS meta-analysis of 5,539 autoantibody positive RA cases and 20,169 controls of European descent, followed by replication in an independent set of 6,768 RA cases and 8,806 controls, which included a total of 8,254,863 SNPs[37]. SLE comprised 7,219 cases and 15,991 controls of European ancestry, yielding a total of 7.915,250 SNPs from a new GWAS, meta-analysis with the published GWAS and a replication study[38]. T1D consisting of 8,781,607 SNPs was extracted from a Mendelian randomization analysis with 5,913 T1D cases and 8,828 reference samples[39]. All the samples in the present study came from Northern and Western European ancestry (CEU) population. The summary statistics have been undertaken genomic control in the individual study level or meta-analysis. Furthermore, the researcher of ImmunoBase applied a global cutoff of MAF<99% for all datasets as determined. The further detailed inclusion criteria and quality control in different GWAS studies were described in the original publications[33–39]. It is worth noting that the summary statistics must include p-values, regression coefficients and standard error after GWAS or meta-analysis.

### Data Preparation

When dealing with the various datasets, we combined the seven diseases’ summary statistics for the common SNPs studied in all datasets firstly. The result of 324,031 overlapping SNPs was selected to perform the multivariate analysis. Second, gene annotation was completed according to the 1000 Genome datasets (website:https://www.cog-genomics.org/static/bin/plink/glist-hg19) using PLINK1.9. Third, the linkage disequilibrium (LD) based SNP pruning method was used to remove SNPs with large pairwise correlations. This SNP pruning method was proceeded by using 50 SNPs as a window where LD was calculated between each pair of SNPs. The minor allele frequency (MAF) was also used as a criterion in the SNP pruning method, which removed the SNP with smaller MAF for pairs with R^2^ > 0.2. Following this initial removal of SNPs in high LD, each sliding window of 5 SNPs forward and the process repeated until there were no pairs of SNPs that are greater than 0.2[40]. All datasets were pruned using the HapMap 3 CEU genotypes as a reference panel (website: http://www.sanger.ac.uk/resources/downloads/human/hapmap3.html) and there remained 41,274 SNPs located in 11,516 gene regions on which we performed the metaCCA analysis. At last, the regression coefficient beta need normalization before conducting the metaCCA analysis if the individual-level data set genotype and phenotype matrices were not standardized. Standardization was achieved afterwards by:

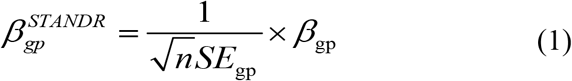

where *SE*_*gp*_ is the standard error of *β*_*gp*_, as given by the original GWAS result, *g* is the number of genotypic, *p* is the number of phenotypic variables, and *n is* the sample number of each diseases.

### MetaCCA Analysis

MetaCCA is an extension of the method of CCA, which requires a full covariance matrix (∑), consisting of four covariance matrices, can be obtained:

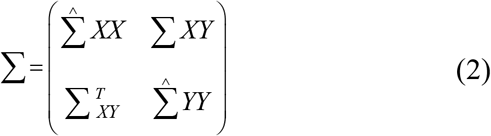

Where ∑^*XY*^ is a cross-covariance matrix between all genotypic and phenotypic variables, 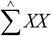 is a genotypic correlation structure between SNPs, 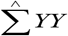 is a phenotypic correlation structure between traits, and 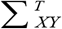 is the transposition of ∑^*XY*^ [16]. ∑^*XY*^ is constructed as the normalized regression coefficient *βgp*:

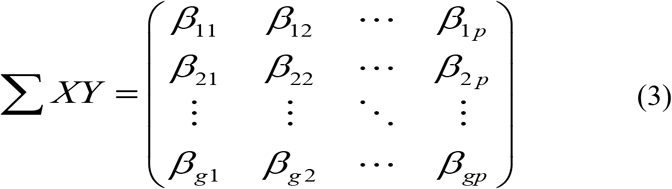

In metaCCA, 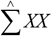 is calculated using a reference database representing the study population, such as the 1000 Genomes database, or other genotypic data available on the target population. There will be better results if 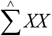 were estimated from the target population or the same ethnicity instead of interracial populations[16]. In our study, 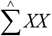 was estimated using the reference SNP dataset of the HapMap 3 CEU.

The phenotypic correlation structure 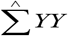 is computed based on ∑^*XY*^. Each entry of 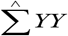 corresponds to a Pearson correlation coefficient between the vector of *β* estimates from *p* phenotypic variables across *g* genetic variants. It has been demonstrated that the bigger *g*, the more accurate the quality of the estimate. Thus, 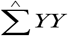 were calculated from summary statistics of 324,031 overlapping SNPs, even if 41,274 of them were used for next analysis.

After calculation, we need determine whether the full covariance matrix is positive semidefinite (PSD). If it is not PSD, an iterative procedure is used to shrink the full covariance matrix until ∑ becomes PSD. In the next analysis, the PSD of the full covariance matrix is plugged into the canonical correlation analysis framework to get the final genotype-phenotype association result[16].The correlation between genotype and phenotype is called the canonical correlation r[41].

In this study, two types of multivariate analysis were considered and the conservative corrected method of Bonferroni was used as the threshold for nominal significance. If the p-value of the canonical correlation r of any SNP was smaller than 1.21×10^−6^ (=0.05/41,274), it was deemed significantly associated with the seven autoimmune diseases. Second, multivariate SNP-multivariate phenotype association analysis at the gene level was tested to identify any potential pleiotropic gene. Similarly, genes with a p-value of canonical correlation smaller than 4.34×10^−6^ (=0.05/11,516) were identified as the potential pleiotropic genes with multiple autoimmune diseases.

### Gene-based association analysis

VEGAS2 (Versatile Gene-based Association Study–2) is a gene-based association method that calculates the correlation analysis of multiple SNPs in a gene region with one phenotype using original GWAS summary statistics[42]. It has been widely applied into the genetic field and shown higher sensitivity and lower false positive rates compared to other gene-based approaches[43]. In the present study, the VEGAS2 method was complementary to the metaCCA and used to refine the identified genes by metaCCA by computing the gene-based p-value for one specific disease. It was performed at: https://vegas2.qimrberghofer.edu.au/, and the threshold was 1.00E-06.

### GO term enrichment analysis

An useful way to understand polygenic associations is to determine whether the implicated genetic variants occur in genes that comprise a biological pathway or not[44]. GO term enrichment analysis classifies gene functions based on three main categories, including molecular function, cellular component and biological process. We conducted the GO term enrichment analysis to find which GO term are over-represented (or under-represented). All significant genes re-identified by VEGAS2 in our study were annotated and enriched (website: http://amp.pharm.mssm.edu/Enrichr/). An adjusted p-value < 0.05 in the enrichment analysis indicates the nominal significance[45].

### Protein-protein interaction network

Protein–protein interaction is crucial for all biological processes and the networks provide many new insights into protein function[46]. In order to detect interactions and associations of all putative pleiotropic genes, the PPI analysis were conducted by searching the Search Tool for the Retrieval of Interacting Genes/Proteins (STRING) database (website: http://string-db.org/), which comprises known and predicted associations from curated databases or high-throughput experiments, also with other associations derived from text mining, co-expression, and protein homology[47].

## Author Contributions

**Conceptualization:** Xiaocan Jia, Xuezhong Shi.

**Data curation:** Xiaocan Jia, YongliYang.

**Formal analysis:** Xiaocan Jia, Nian Shi, Yifan Li, Jiebing Tan.

**Methodology:** Xiaocan Jia, Nian Shi, Zhenhua Xia, Yu Feng.

**Funding acquisition:** wei Wang, Changqing Sun, Hongwen Deng

**Project administration:** Xuezhong Shi, Yongli Yang.

**Software:** Zhenhua Xia, Yu Feng, Fei Xu.

**Supervision:** Xuezhong Shi, Wei Wang.

**Writing – original draft:** Xiaocan Jia, Xuezhong Shi.

**Writing – Review & Editing:** Nian Shi, Yongli Yang, Changqing Sun.

